# SETDB1 promotes tubulin deacetylation and Golgi fragmentation by HDAC6

**DOI:** 10.64898/2026.06.24.734187

**Authors:** Gowthaman Gunasekaran, Galia Gelman, Ohad Manshirov, Tamar Listovsky, Gabi Gerlitz

## Abstract

Microtubules (MTs) are dynamic cytoskeletal structures essential for intracellular transport, cell division, and organelle positioning. Their functions are regulated by post-translational modifications, including α-tubulin acetylation at Lys40, which enhances MT stability and resilience. Histone deacetylase 6 (HDAC6) is the primary enzyme that reverses this modification, but its access to the luminal Lys40 residue is restricted. Previously, we identified SETDB1, a histone methyltransferase and known oncogene, as a cytoplasmic regulator of MT dynamics, attenuating MT polymerization and destabilizing MTs. Here, we uncover the molecular mechanism by which SETDB1 destabilizes MTs. SETDB1 interacts with HDAC6 and promotes its tubulin deacetylation activity. Mechanistically, SETDB1 enhances HDAC6 recruitment to polymerized MTs and induces repairable damage along MT shafts, generating entry points for HDAC6 into the MT lumen. Functionally, this axis regulates Golgi organization: SETDB1 overexpression disperses the Golgi in an HDAC6-dependent manner, while SETDB1 knockdown or HDAC6 inhibition compacts it. Notably, SETDB1’s role in Golgi regulation is independent of its methyltransferase activity. These findings reveal crosstalk among the histone methylation machinery, MT dynamics, and Golgi organization. Since Golgi dispersal is thought to promote tumorigenesis, our results suggest that the SETDB1-HDAC6 axis is a potential therapeutic target.

**Graphical Abstract:** 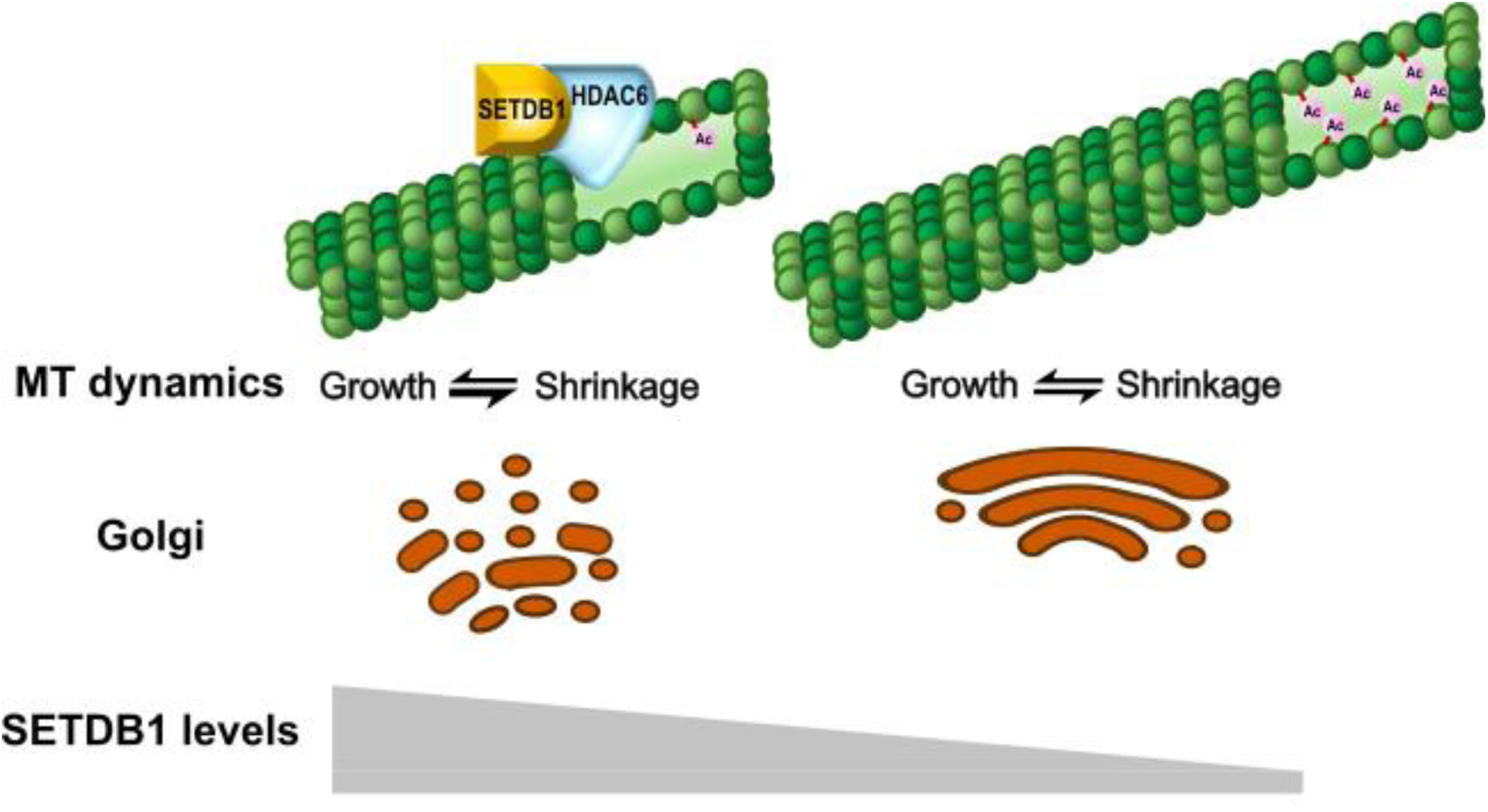

## Introduction

Microtubules (MTs) are a fundamental part of the cytoskeleton involved in different cellular functions, including cellular shape determination, chromosome segregation in mitosis, nucleokinesis, differentiation, intracellular cargo transportation, and the regulation of structure and localization of membrane-bound organelles [1–6]. MTs are composed of α- and β-tubulin heterodimers that undergo cycles of growth and shrinkage. This dynamic instability supports the various cellular functions of the MTs [7]. MT functions can be modulated by post-translational modifications (PTMs), including acetylation, detyrosination, phosphorylation, glutamylation, and lactylation. MT PTMs generate a “tubulin code” that modulates MT mechanical properties and interactions with microtubule-associating proteins (MAPs) [8–10]. Acetylation at α-tubulin Lys40 (acetylated tubulin) in the lumen of the MT reduces lateral contacts between MT protofilaments and increases MT flexibility, making MTs more resilient to mechanical aging, less prone to fragmentation, and more stable [11,12]. This acetylation is mediated by the forward enzyme *N*-acetyltransferase 1 (ATAT1) and reversed mainly by histone deacetylase 6 (HDAC6) and less often by sirtuin 2 (SIRT2) [13–15]. Whereas ATAT1 can penetrate the lumen from the ends of the MT cylinder or through gaps between protofilaments [16–18], HDAC6 penetration from the cylinder ends is much less efficient; thus, it is believed that it prefers tubulin dimers as substrates [19,20]. Through tubulin deacetylation, HDAC6 affects various processes, including migration, endocytosis, proliferation, and mitosis [21,22].

Recent studies identified moonlight functions for several factors involved in histone methylation in the regulation of MT dynamics. The histone methyltransferase SETD2 binds α-tubulin to generate αTubK40me3 during mitosis to support mitotic fidelity and to prevent genomic instability [23–26]. In neurons, αTubK40me3 by SETD2 promotes MT polymerization to support neuronal migration of cortical neurons during embryogenesis [27]. In hippocampal neurons, SETD2 supports dendritic arborization and axon extension along with prevention of anxiety in mice [28]. Moreover, αTubK40me3 is read by PBRM1, which is a subunit of the chromatin remodeling complex PBAF [29], and is erased by KDM4A [30]. Additionally, the histone methyltransferase SET8 binds MTs and methylates α-tubulin at Lys311 [31]. The histone methyltransferase MLL1 with its co-factor WDR5 promote the localization of kinesin Kif2A to spindle MTs to support mitotic fidelity [32]. MLL1/WDR5 also localize to the pericentriolar material and interact with centriolar satellite protein Cep72 and γ-tubulin ring complex proteins (γ-TuRCs) to promote MT polymerization and spindle formation [33].

An additional histone methyltransferase involved in MT regulation is SETDB1. SETDB1 methylates histone H3 at Lys9 to induce heterochromatin formation and gene repression. SETDB1 is considered an oncogene that is amplified in various cancers, including gynecologic, breast, lung, and skin cancers, and its oncogenic behavior supports cancer cell proliferation, escaping from the immune response, migration, and invasion [34–37]. SETDB1 shuttles between the nucleus and the cytoplasm due to its nuclear localization signal (NLS) and nuclear export signal (NES) motifs [38,39]. While SETDB1’s nuclear roles are well-documented, its cytoplasmic functions remain poorly understood. In embryonic stem cells, cytoplasmic SETDB1 was found to interact with the RNA-binding protein Trim71 to support its activity in regulation of mRNA metabolism and translation [40]. In cancer cells, we found that cytoplasmic SETDB1 interacts with MTs and attenuates their polymerization. In support of it, silencing of SETDB1 led to increased duration of mitosis and mitotic catastrophes in HeLa cells. Mechanistically, the ability of SETDB1 to attenuate MT polymerization is maintained in a catalytic-dead version of the protein, suggesting it is methyltransferase-independent. We identified an association of SETDB1 with HDAC6, along with increased tubulin acetylation in SETDB1 KD cells [41].

However, the molecular basis of how SETDB1 facilitates HDAC6-mediated tubulin deacetylation remains elusive, as well as the cellular consequences of this regulatory mechanism. Here, we found that SETDB1 interacts with HDAC6 and functions as a cofactor to enhance HDAC6-mediated tubulin deacetylation. SETDB1 promotes HDAC6 association with polymerized MTs rather than soluble tubulin dimers, and it promotes MT shaft damage, which could serve as penetration sites of HDAC6 into the MT lumen. Importantly, SETDB1-dependent activation of HDAC6 is involved in determining Golgi organization, supporting a functional link between MT acetylation and organelle organization.

## Materials and Methods

### Cell Culture

Cells were maintained in Dulbecco’s Modified Eagle Medium (DMEM; 3202828, Gibco, Thermo Fisher Scientific, Paisley, UK) supplemented with 10% fetal bovine serum (FBS), 1% L-glutamine, and 0.5% penicillin-streptomycin. Cells were cultured at 37°C in a humidified incubator with 7% CO_2_. Plasmid-DNA transfections were performed using jetOPTIMUS DNA transfection reagent (0000004435, Polyplus Illkirch-Graffenstaden, France) according to the manufacturer’s instructions. Cells were incubated for 24h at 37°C post transfection prior to further analysis. For gene knockdown, cells were transfected with siRNA (IDT, Coralville, IA, USA) using INTERFRin transfection reagent (0000004693, SARTORIUS, Göttingen, Germany) according to the manufacturer’s instructions. siRNAs used were human SETDB1 (hs.Ri.SETDB1.13.1) and negative control (51-01-14-04). Cells were incubated for 72h at 37°C post transfection prior to further analysis.

### Co-Immunoprecipitation

HEK293 cells were co-transfected with HDAC6-FLAG (a gift from Tso-Pang Yao, Addgene plasmid # 30482) [42] and GFP-SETDB1 [41] or pEGFPC3 (Clontech, TaKaRa) and incubated at 37°C for 24h, in a 7% CO_2_ environment. Cells were lysed in extraction buffer [43] for 30min on ice. Lysates were clarified by centrifugation at 20,000g for 20min at 4°C. A 5% input sample was collected from clarified lysate and denatured in SDS sample buffer by boiling at 95°C for 10min. For immunoprecipitation, clarified lysates were incubated with anti-α-FLAG antibody (CST-14793S, Cell Signalling Technology, Danvers, Massachusetts, USA) pre-bound to protein A/G magnetic beads (LSKMAGAG02, Millipore, USA) on a rotor for 1h at 4°C. Beads were washed three times with extraction buffer and once with cold PBS, then boiled in SDS sample buffer at 95°C for 10min before separation on SDS-PAGE for Western blot analysis.

### Co-sedimentation assay

Cells were co-transfected with HDAC6-FLAG and GFP-SETDB1 or pEGFPC3 and incubated at 37°C and 7% CO_2_ for 24h. Cells were lysed in PIPES-buffer (80 mM PIPES, pH 6.8, 1 mM MgCl_2_, 1 mM EGTA, 100 mM NaCl, 1% Triton X-100) supplemented with 1x protease inhibitor cocktail (K1007, APExBIO, Houston, Texas, USA) for 30 minutes on ice. Lysates were clarified by two repetitive centrifugation steps at 20,000g for 20min at 4°C. Supernatants were supplemented with 1mM GTP (22625, ThermoFisher Scientific, China) and 40μM paclitaxel (32842, ThermoFisher Scientific, USA) and incubated at 4°C or 37°C. Samples were centrifuged at 20,000g at 4°C or 37°C, respectively. The resulting supernatant and pellets were subjected to Western blot analysis.

### Microtubule damage staining

To visualize the microtubule shaft damage, HeLa cells were transfected with GFP-SETDB1 [41] or GFP-Radixin [44] GTP-Tubulin staining was carried out as reported previously [45,46]. Briefly, cells were permeabilized with GPEM buffer (80mM PIPES pH 6.9, 2mM EGTA, 1mM MgCl_2_ and 0.2% glycerol) for 3min at 37°C followed by anti-tubulin-GTP (hMB11, 1:10,000, AG-27B-0009, Adipogen, San Diego, USA) incubation at 37°C for 10min. After 2 washes with prewarmed GPEM buffer, cells were incubated with anti-human-Cy3 antibody (171258, Jackson Immunoresearch Laboratories, Pennsylvania, USA) for 15min at 37°C. After one wash with prewarmed GPEM buffer, cells were fixed with methanol supplemented with 1mM EGTA at -20°C for 6min and stained with goat anti-GFP (1:300, GTX26673, GeneTex, Irvine, California, USA), mouse anti-α-tubulin (1:200, DM1A, sc-32293, Santa Cruz, Texas, USA) antibodies, and Hoechst 33342 (B2261, Sigma-Aldrich, Rehovot, Israel). Images were collected using an Olympus IX81 fluorescence microscope with a coolSNAP HQ2 CCD camera (Photometrics, Tucson, AZ, USA) or a Prime BSI Express camera (Teledyne Photometrics, Tucson, AZ, USA).

### Immunofluorescence

Cells were plated on coverslips and transfected with the indicated plasmid DNA or siRNA. After 24h of plasmid DNA or 72h of siRNA transfection and treated with 1.5 µM Tubastatin A (15785, Cayman Chemical, Ann Arbor, Michigan, USA) for 1h at 37°C. Cells were fixed with 3% paraformaldehyde (PFA) for 10min at room temperature. Antibodies used included goat anti-GFP (1:300, GTX26673, GeneTex, Irvine, California, USA), rabbit anti-GM130 (1:500, CST12480, Cell Signaling Technology, Danvers, Massachusetts, USA), mouse anti-α-tubulin (1:200, DM1A, sc-32293, Santa Cruz, Texas, USA) and Hoechst 33342 (B2261, Sigma-Aldrich, Rehovot, Israel). Microscopy images were captured using an Olympus 1X81 fluorescent microscope equipped with a coolSNAP HQ2 CCD camera (Photometrics, Tucson, AZ, USA) or a Prime BSI Express camera (Teledyne Photometrics, Tucson, AZ, USA). Quantification was done by the ImageJ/Fiji software (National Institutes of Health, Bethesda, United States).

### In vitro deacetylation assay

HEK293 cells were plated on 6cm culture dishes and transfected with GFP-SETDB1 or pEGFPC3. Following an incubation for 24h, cells were harvested with cold PBS and centrifuged at 500g for 5min at 4°C. Lysed with extraction buffer for 30min on ice and centrifuged at 20,000g for 20min at 4°C. A 5% input sample was collected from clarified lysate and denatured in SDS sample buffer by boiling at 95°C for 10min. For pull-down, clarified lysates were added to GFP-selector magnetic agarose beads (N0315, NanoTag Biotechnologies, Germany) and incubated on a rotor for 1h at 4°C. The beads were washed thrice with extraction buffer (supplemented with 0.05% BSA) and once with cold PBS. The extracted beads were resuspended in BRB80 buffer (80mM PIPES pH 6.9, 1mM EGTA and 1mM MgCl_2_) [47,48] for the *in vitro* assay.

For the *in vitro* assay, 0.4μM acetylated tubulin dimers (CS-TAC01, Cytoskeleton, Inc, Denver, Colorado, USA), 0.5nM recombinant HDAC6-FLAG enzymes (50056, BPS Bioscience, San Diego, USA) were added to the BRB80 buffer. GFP pull-down samples containing GFP-SETDB1 or GFP proteins were added to the reactions. The deacetylation reaction was carried out at 37°C for 1h and terminated by the addition of 5X SDS sample buffer followed by boiling at 95°C for 10min. Samples were further subjected to Western blot analysis.

### SDS-PAGE and Western blot analysis

Cells were harvested, washed in PBS buffer, and centrifuged at 500g for 5min at 4°C. Lysates were prepared by sonication in 2X SDS sample buffer (100 mM Tris 6.8, 2% SDS, 10% glycerol, 100 mM DTT) supplemented with 1X protease inhibitor cocktail (K1007, APExBIO, Houston, Texas, USA) followed by heating at 95°C for 10 min. The proteins were separated by SDS–PAGE and transferred to nitrocellulose membranes. The following antibodies used for western blotting: rabbit anti-SETDB1(1:500, GTX115305, GeneTex, Irvine, California, USA), rabbit anti-HDAC6 (1:10,000, ab133493, Abcam, Cambridge, UK), mouse anti-α-tubulin (1:500, DM1A, sc-32293, Santa Cruz, Texas, USA), rabbit anti-acetylated α-tubulin (1:1000, CST5335, Cell Signaling Technologies, Danvers, Massachusetts, USA), rabbit anti-FLAG (1:1000, CST14793, Cell Signalling Technology, Danvers, Massachusetts, USA), and mouse anti-β-actin (1:500, sc-69879, Santa Cruz, Texas, USA). Detection was performed using appropriate secondary antibodies. Quantification was done by the ImageJ/Fiji software (National Institutes of Health, Bethesda, United States).

## Results

### SETDB1 promotes tubulin deacetylation

SETDB1 association with the main tubulin deacetylase HDAC6 was demonstrated previously by co-immunoprecipitation of overexpressed proteins [41]. To validate this interaction under more physiological conditions, we performed co-immunoprecipitation of endogenous proteins in HEK293 cells. As shown in **Figure 1** and **Sup. Figure 1**, immunoprecipitation of SETDB1 resulted in co-pull down of endogenous HDAC6, confirming the interaction at endogenous expression levels.

**Figure 1.**
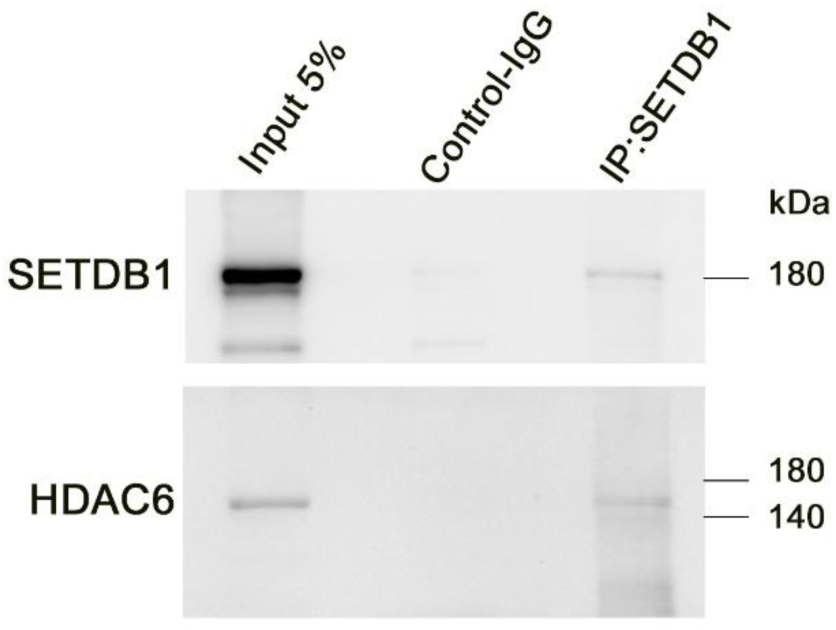
Co-immunoprecipitation of endogenous SETDB1 and HDAC6 in HEK293 cells. HEK293 cell lysates were immunoprecipitated with control rabbit IgG or an anti-SETDB1 antibody (CST-2196) and analyzed by Western blot for the indicated proteins.

Previously, we found that SETDB1 knockdown led to elevated tubulin acetylation levels in mouse melanoma cells [41]. To further establish the ability of SETDB1 to promote tubulin deacetylation we evaluated the effect of SETDB1 overexpression on tubulin acetylation in human melanoma cells WM266.4. The basal tubulin acetylation level in these cells is very low; therefore, we treated the cells with the MT stabilizing drug paclitaxel. Paclitaxel as other taxanes is known to promote tubulin acetylation [49,50], thus it can generate a substantial pool of acetylated tubulin. As expected, paclitaxel treatment increased the acetylated tubulin levels in a dose-dependent manner (**Figure 2**). Notably, overexpression of SETDB1 consistently reduced tubulin acetylation levels to 27-48% of the levels observed in GFP expressing control cells, supporting the notion that SETDB1 is a positive co-factor of HDAC6.

**Figure 2.**
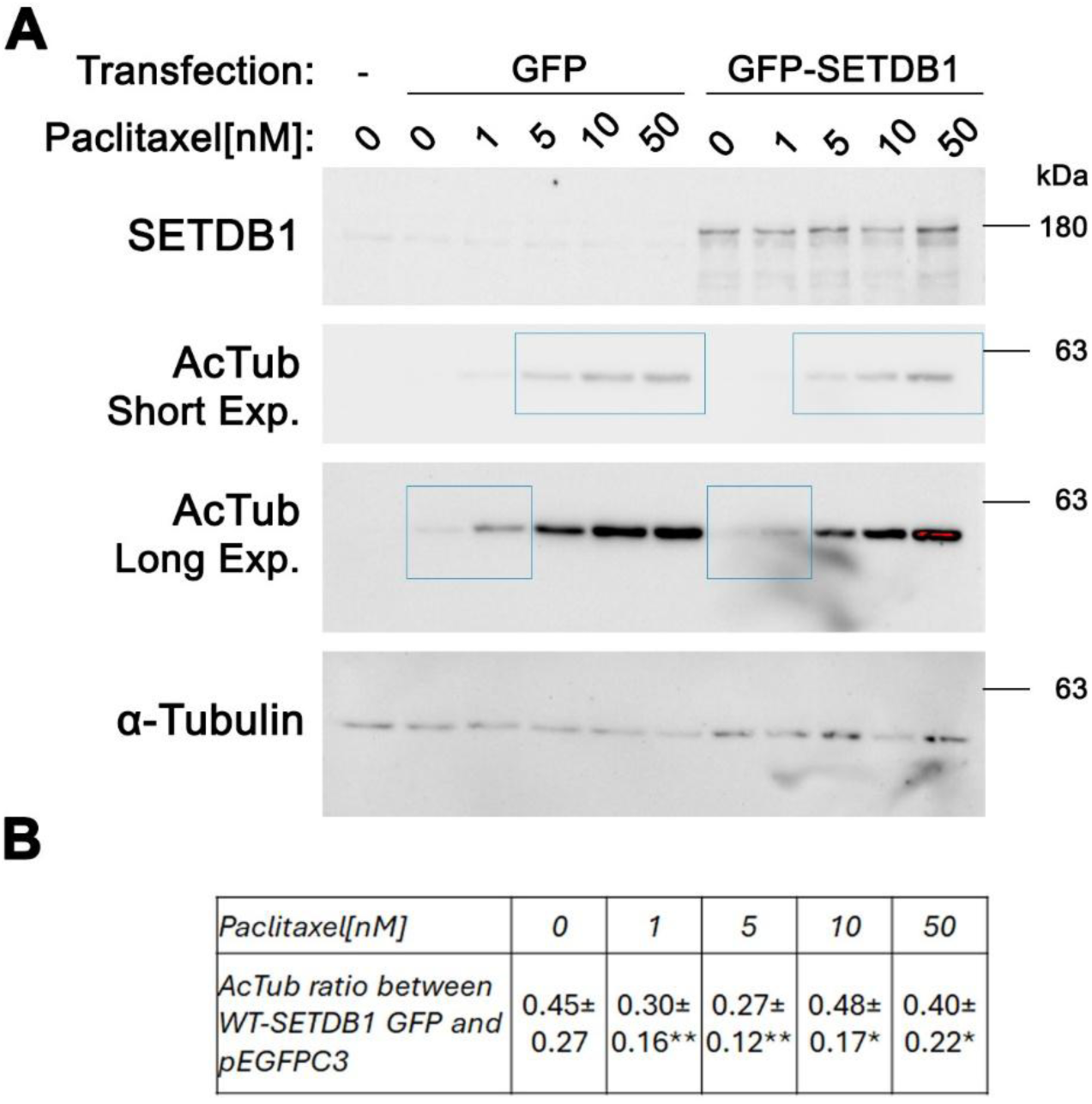
SETDB1 overexpression reduced acetylated tubulin levels. **A.** WM266.4 cells expressing GFP or GFP-SETDB1 treated with the indicated paclitaxel concentrations for 1 h and subjected to Western blot analysis for the indicated proteins. Due to the wide dynamic range of acetylated tubulin (AcTub) levels, both short and long exposures are shown. Blue rectangles indicate the lanes used for AcTub quantification. **B.** Ratios of AcTub levels in GFP-SETDB1-expressing cells relative to GFP-expressing control cells. AcTub signals were normalized to total α-tubulin levels, and the ratios of AcTub in GFP-SETDB1-expressing cells relative to GFP-expressing cells were calculated. Data are represented as the mean ± SE of three independent experiments. Statistical significance was evaluated by the Student’s *t*-test, \**p* < 0.05, \*\**p* < 0.01.

To determine if SETDB1 can promote directly HDAC6 enzymatic activity, we added immunoprecipitated SETDB1 to an *in vitro* tubulin deacetylation reaction containing recombinant HDAC6. As shown in **Figure 3**, the presence of SETDB1 increased HDAC6-dependent tubulin deacetylation by approximately 25%, indicating that SETDB1 acts as a positive regulator of HDAC6 activity.

**Figure 3.**
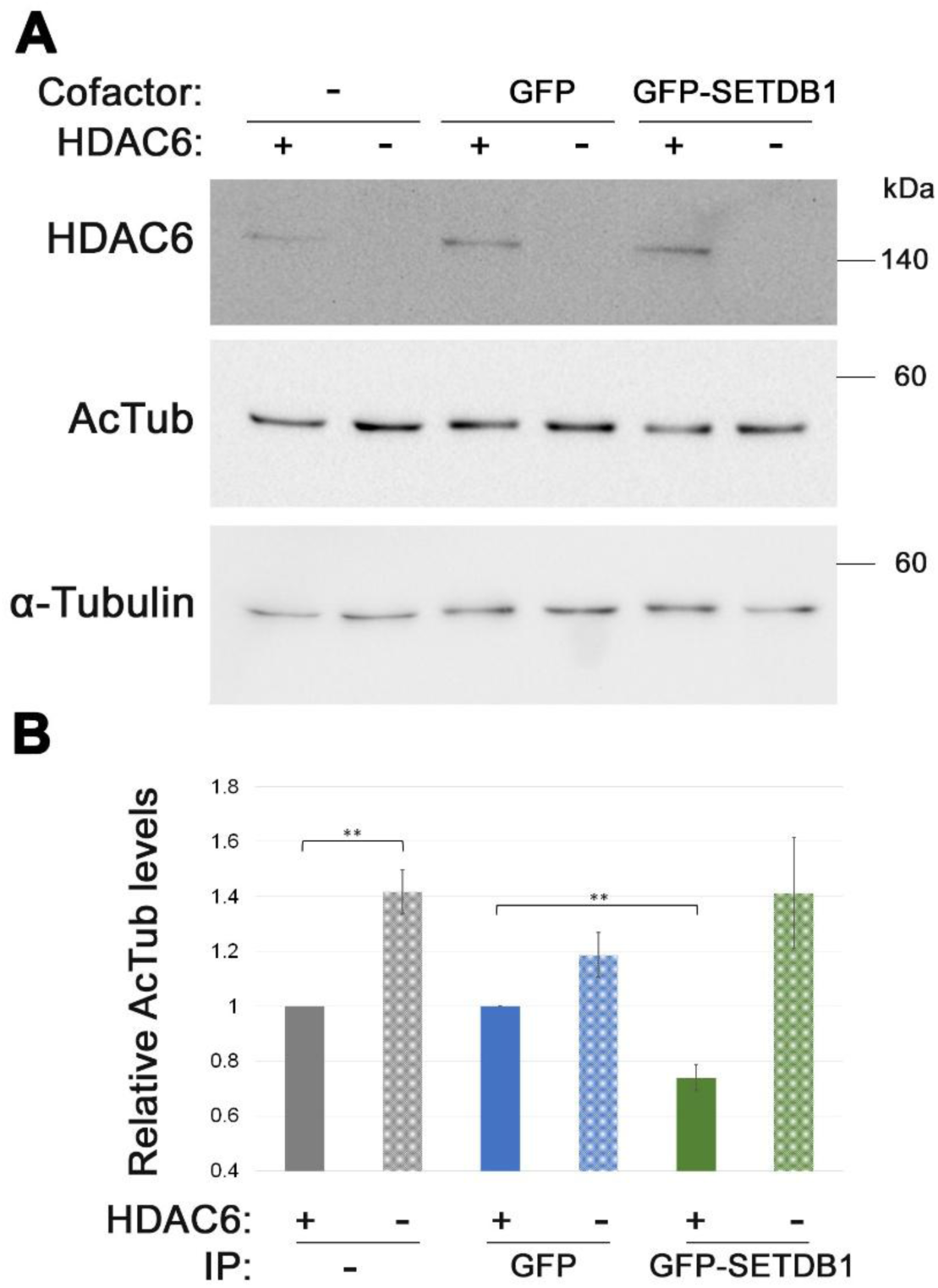
SETDB1 affects HDAC6 activity *in vitro*. **A.** *In vitro* deacetylation of α-tubulin by recombinant HDAC6, alone or with immunoprecipitated GFP or GFP-SETDB1 (cofactor), was analyzed by Western blot. **B.** Bar graph of mean acetylated tubulin (AcTub) levels ± SE of three independent repetitions. AcTub signals were first normalized to total α-tubulin levels. For samples without additional cofactor the HDAC6-only sample was set as 1. For samples with immunoprecipitation (IP), the HDAC6+ GFP sample was set as 1. Statistical significance was calculated with Student’s *t*-test, \*\**p* < 0.01.

### SETDB1 promotes HDAC6 association with MTs

To gain insight into how SETDB1 enhances HDAC6 function, we examined the ability of HDAC6 to bind tubulin dimers in the presence of SETDB1. While immunoprecipitation of overexpressed HDAC6 co-pulled tubulin dimers, co-expression of SETDB1 unexpectedly reduced this association (**Figure 4**). This observation aligns with our previous findings that SETDB1 was not co-immunoprecipitated with soluble tubulin dimers (unshown data) but did co-sediment with polymerized MTs [41]. We therefore assessed whether SETDB1 affects HDAC6 binding to MTs. Indeed, SETDB1 overexpression increased the association of HDAC6 with MTs by ∼1.4-fold in HEK293 cells (**Sup. Figure 2**) and ∼3-fold in WM266.4 cells (**Figure 5**), demonstrating a SETDB1-dependent enhancement of HDAC6 recruitment to the MT network.

**Figure 4.**
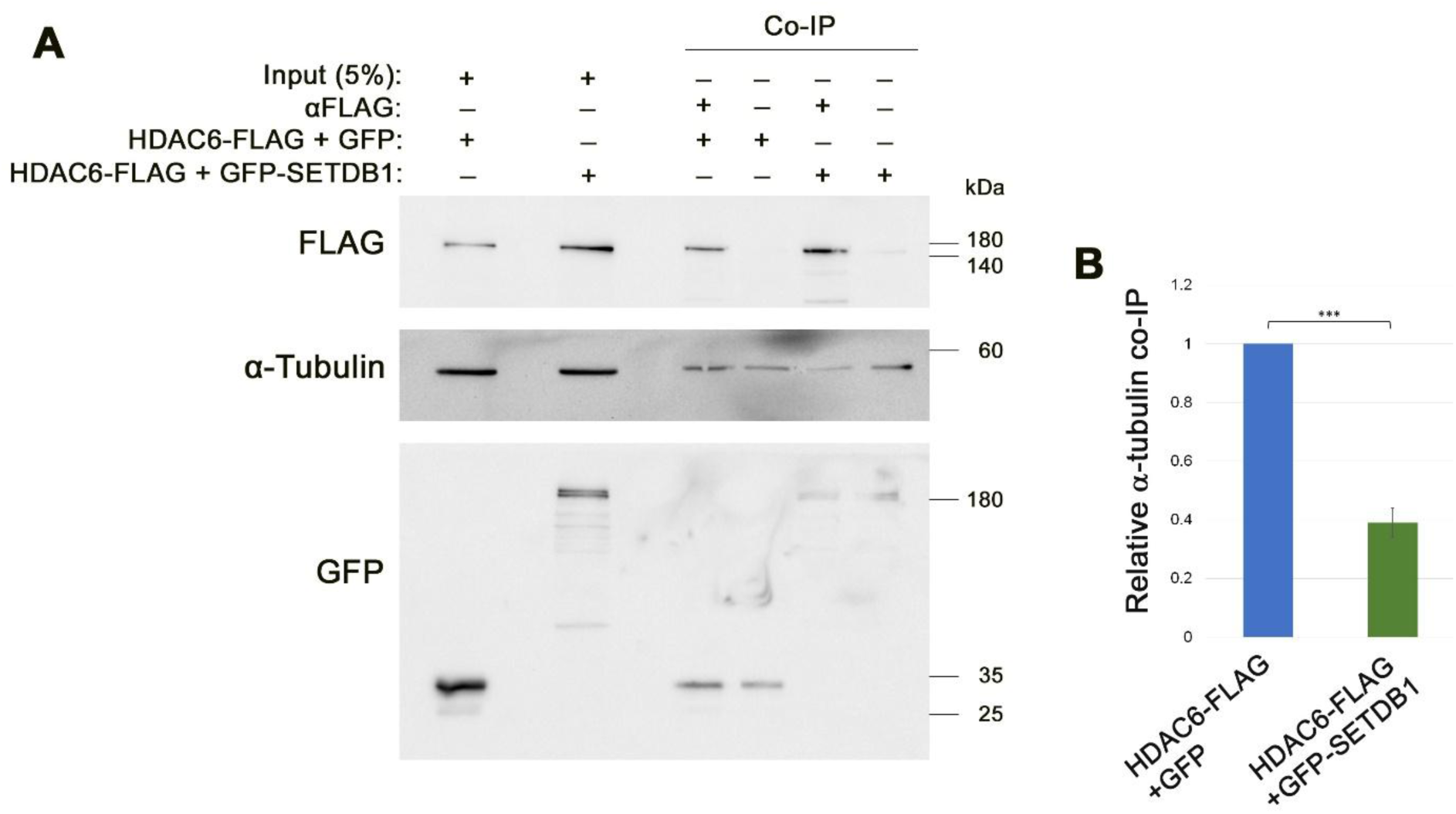
SETDB1 reduced HDAC6 co-immunoprecipitation with α-tubulin. **A.** HEK293 cells co-expressing HDAC6-FLAG with GFP or GFP-SETDB1 were immunoprecipitated with anti-FLAG antibodies and analysed by Western blot for the indicated proteins. **B.** Bar graph showing the mean relative levels of α-tubulin co-immunoprecitated with HDAC6-FLAG ± SE of three independent repetitions. In each repetition, co-immunoprecipitated α-tubulin levels were normalized to HDAC6-FLAG pull down levels and the ratio in GFP-expressing cells was set to 1. Statistical significance was calculated with Student’s *t*-test, \*\*\**p* < 0.001.

**Figure 5.**
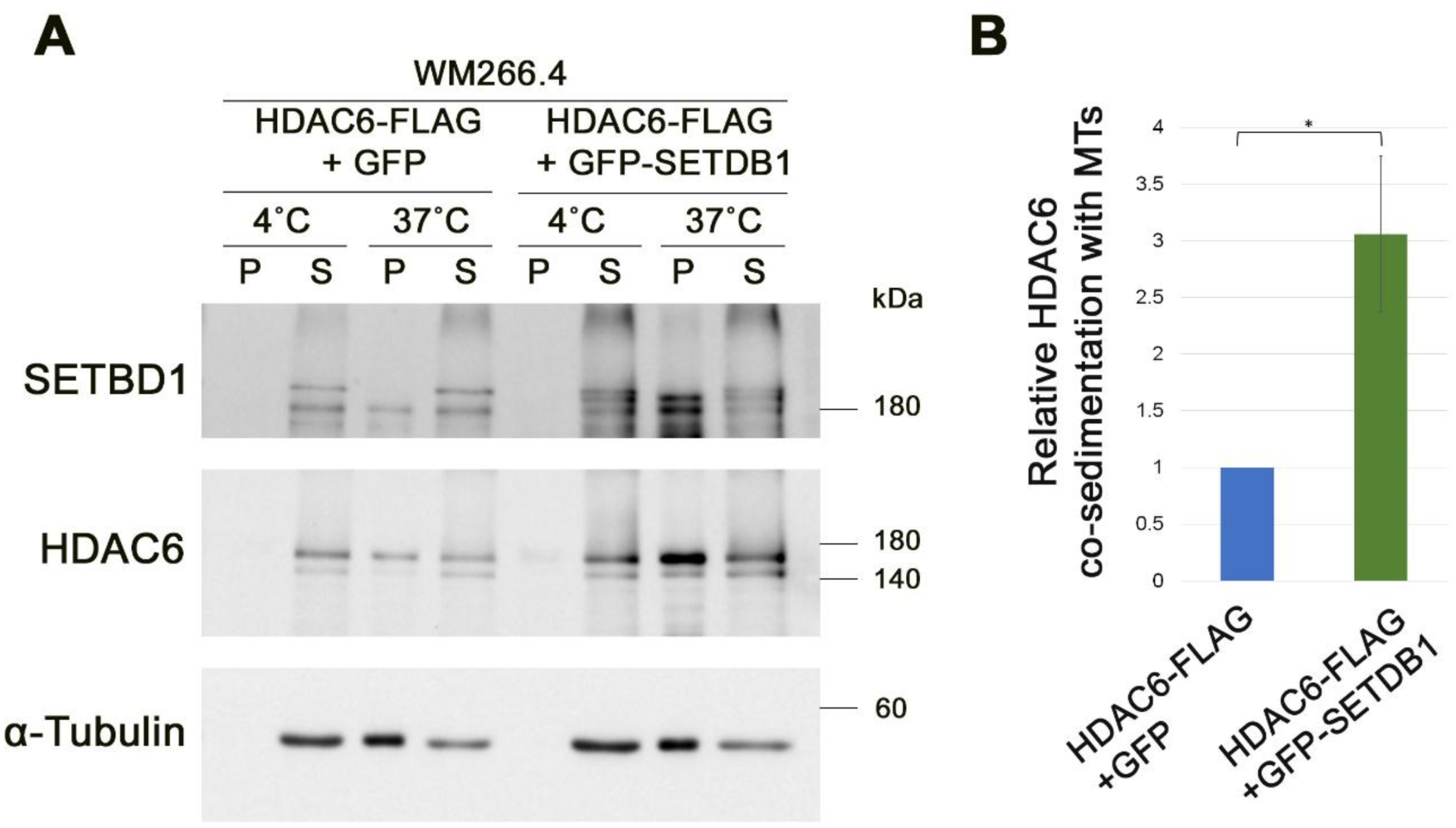
SETDB1 promotes HDAC6 co-sedimentation with MTs. **A.** MT co-sedimentation assay of HDAC6 in WM266.4 cells. WM266.4 cells overexpressing HDAC6-FLAG with GFP or GFP-SETDB1 used for MT co-sedimentation assay. The pellet (P) and supernatant (S) obtained after incubation at 37°C indicate MT-bound and unbound fractions, respectively. Incubation at 4°C serves as the MT-free control. **B.** The bar graphs show the relative levels of HDAC6 associated with MTs ± SE of four independent repetitions. In each repetition HDAC6 levels in the pellet were normalized to its levels at the supernatant at 37°C, and the ratio in GFP expressing cells was set to 1. Statistical significance was calculated with Student’s *t*-test, \**p* < 0.05.

The substrate of HDAC6, Lys40 of α-tubulin faces the lumen of the MTs [51]. Thus, improved recruitment of HDAC6 to the surface of MTs is not sufficient to promote its activity. Moreover, whereas the tubulin acetyltransferase ATAT1 penetrates the microtubule lumen with high efficiency [16,18,52], HDAC6 is thought to do so much less effectively [10,19,53,54]. Recently, kinesin-1 was shown to facilitate HDAC6 activity towards MTs by inducing damage along the MT shaft that can be used as penetration points for HDAC6 into the MT lumen [55]. This led us to the hypothesis that SETDB1 may similarly promote limited, repairable damage along MT shafts to generate entry points for HDAC6. Damage along MT shafts leads to repair by the incorporation of GTP-αTubulin that can be detected by the hMB11 antibody [45]. Using hMB11 antibody to detect these exchange sites revealed a 2-fold increase in hMB11 staining in HeLa cells overexpressing SETDB1 in comparison to Radixin-expressing control cells (**Figure 6**). Together, these findings suggest that SETDB1 promotes the association of HDAC6 with MTs along with generation of penetration sites for HDAC6 into the MT lumen.

**Figure 6.**
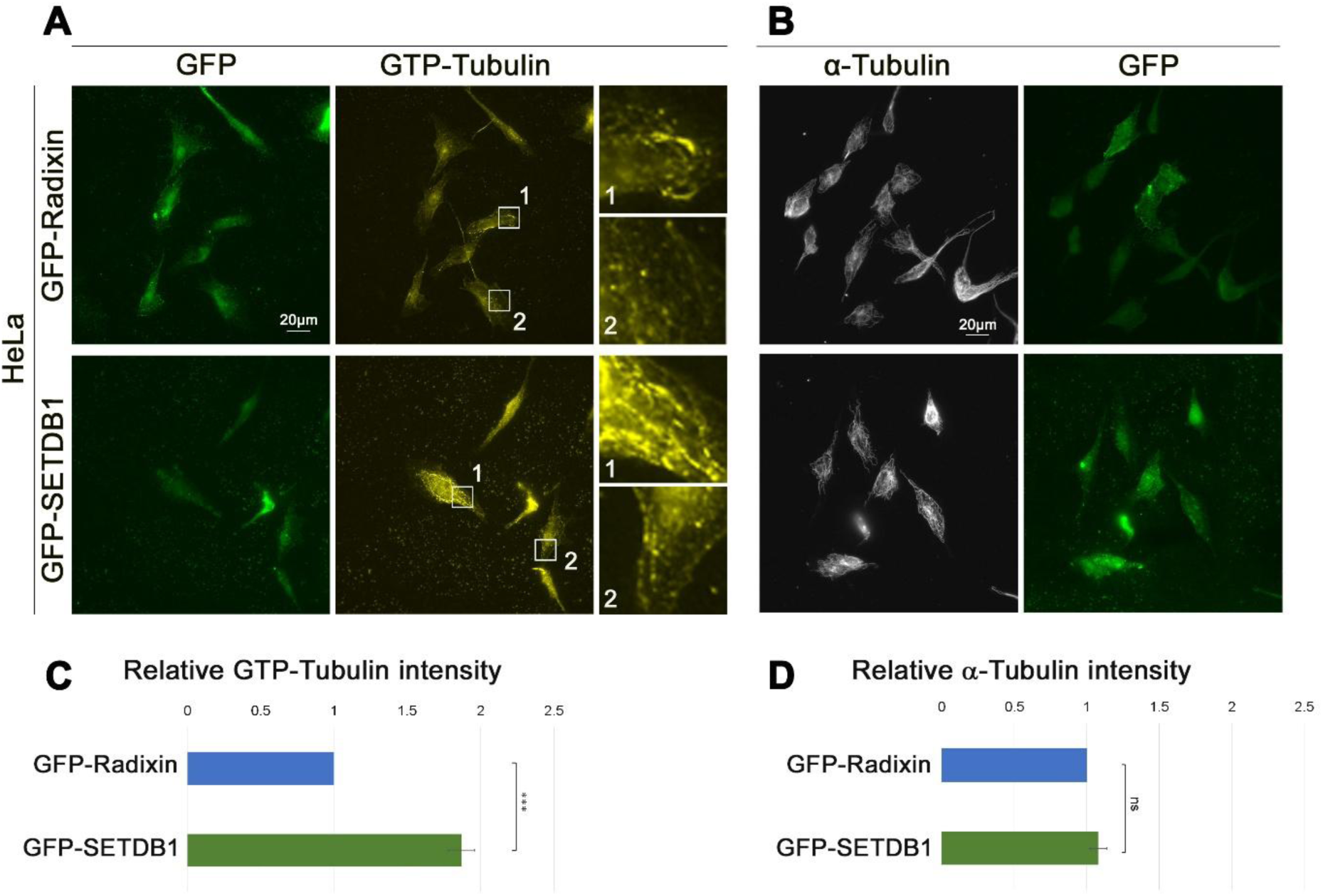
MT shaft damage induced by SETDB1. **A-B.** GFP-SETDB1 and GFP-Radixin expressing HeLa cells immunostained for GTP-tubulin (GTP-Tub) and GFP (A) or α-tubulin and GFP (B). Scale bar: 20 μm. **C-D.** Quantification of GTP-Tub (C) and α-tubulin (D) immunostaining intensities. For each experiment, 18-49 cells per transfection were measured for mean cellular intensity. The average mean intensity was normalized to GFP-Radixin expressing cells. The average mean intensities from three independent experiments ± SE are presented. Statistical significance was calculated with Student’s t-test, \*\*\**p* < 0.001.

### SETDB1 affects Golgi organization in an HDAC6-dependent manner

To examine the physiological consequences of SETDB1-mediated HDAC6 activation, we assessed their combined roles in regulating Golgi morphology. Depolymerization of MTs or inhibition of the dynein motor were shown to induce dispersal of the Golgi apparatus [56–58]. Since acetylated MTs generate a preferred track for dynein [59], we anticipated that interference with tubulin acetylation by overexpression of SETDB1 might compromise Golgi compaction.

Using GM130 immunostaining, we evaluated Golgi morphology while using Dynamitin overexpression as a positive control for Golgi dispersal. Dynamitin is a subunit of the dynein co-factor Dynactin that serves as a dominant negative form of dynein upon overexpression [56,60]. As expected, Dynamitin overexpression led to significant Golgi fragmentation as was evident from increased number of Golgi vesicles per cell along with reduced area of each vesicle (**Figure 7**). Interestingly, overexpression of SETDB1 also increased the dispersal of the Golgi to ∼4.5 times more Golgi vesicles per cell. Inhibition of HDAC6 with Tubastatin A (TubA) interfered with the ability of SETDB1 to disperse the Golgi, indicating that SETDB1 relies on HDAC6 activity to modulate the Golgi organization. Previously we found that methyltransferase-dead mutant SETDB1 altered MT polymerization to the same extent as wildtype SETDB1 [41]. Also in the current assay, catalytic dead (CD) version of SETDB1 affected the morphology of the Golgi as the wildtype protein, suggesting that SETDB1’s role in Golgi regulation does not require its catalytic activity.

**Figure 7.**
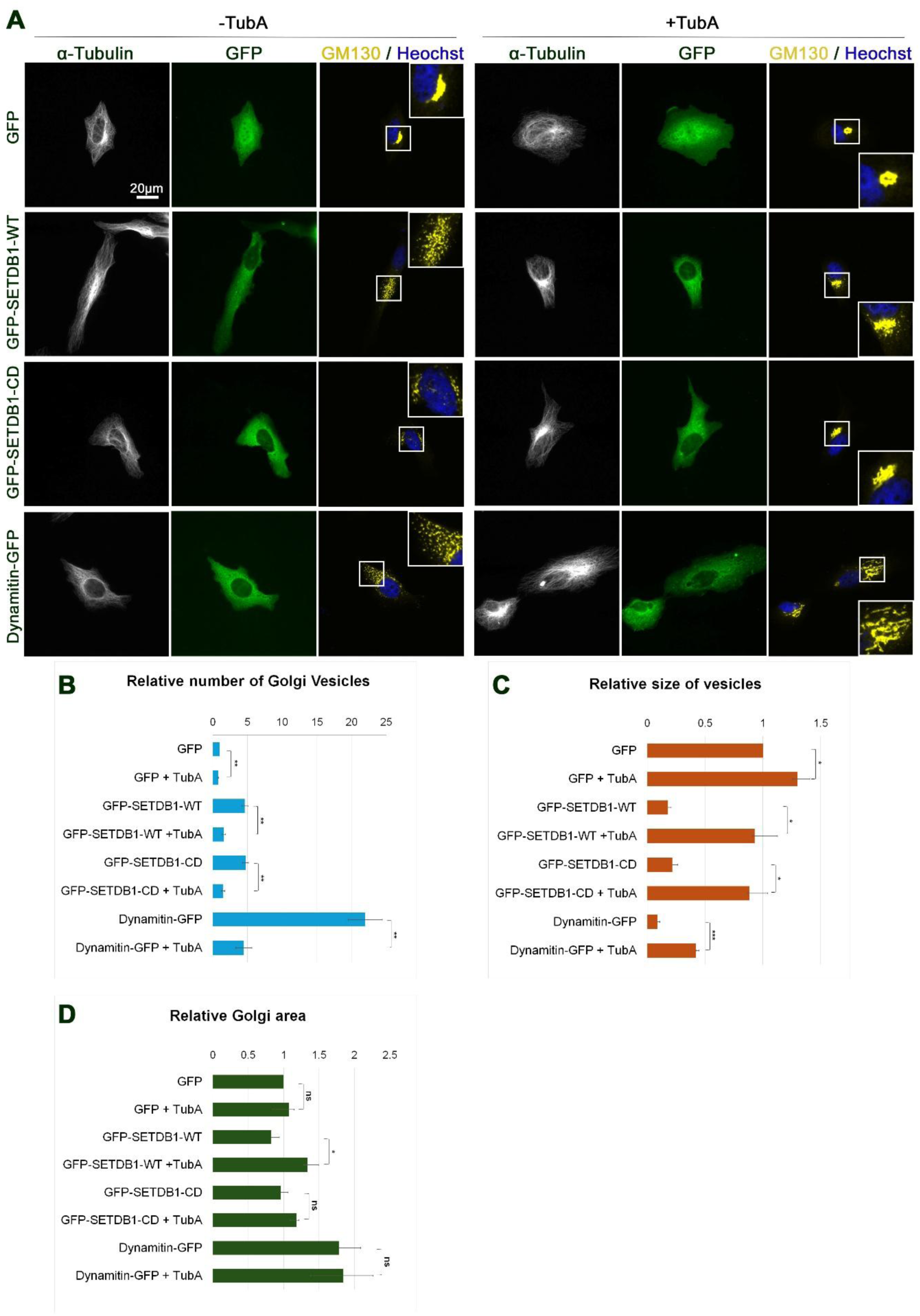
Overexpression of SETDB1 induces Golgi fragmentation. **A.** HeLa cells expressing GFP, GFP-SETDB1 wildtype (WT), GFP-SETDB1 catalytic dead (CD) and Dynamitin-GFP treated or non-treated for 1 h with 1.5 µM of the HDAC6 inhibitor Tubastatin A (TubA) immunostained for α-tubulin, GFP and the Golgi marker GM130. DNA was stained with Hoechst 33342. The Golgi outlined by white rectangles are magnified at the side of the micrographs. Scale bar: 20 μm. **B-D.** Quantification of the relative Golgi dispersal. For each experiment, 21 cells per condition were measured by ImageJ using analyze particle tool for the number of Golgi vesicles per cell (B), average size of Golgi vesicle per cell (C) and total Golgi area (D). In each experiment the averages in each condition were normalized to TubA-free GFP-transfected cells. Average mean intensities from three independent experiments ± SE are presented. Statistical significance was calculated with Student’s t-test, \**p* < 0.05, \*\**p* < 0.01, \*\*\**p* < 0.001.

Since overexpression of SETDB1 led to Golgi dispersal, we wanted to evaluate if SETDB1 knockdown (**Sup. Figure 3**) would lead to accumulation of Golgi vesicles. Towards this aim, we took advantage of the frequent Golgi dispersal in tumor cells. In both human osteosarcoma cells, U2OS and human melanoma cells, WM266.4 the Golgi morphology was dispersed (**Figure 8**). Notably, either SETDB1 KD or HDAC6 inhibition by TubA led to reduction in the number of Golgi vesicles per cell in parallel with an increase in the size of the Golgi vesicles. Thus, suggesting that upon interference with the SETDB1-HDAC6 axis, the Golgi vesicles better fuse to each other to form structures closer to classical Golgi sacs. These results suggest that SETDB1-driven MT deacetylation disrupts dynein-dependent Golgi compaction, a process that could affect protein secretion and glycosylation in cancer cells.

**Figure 8.**
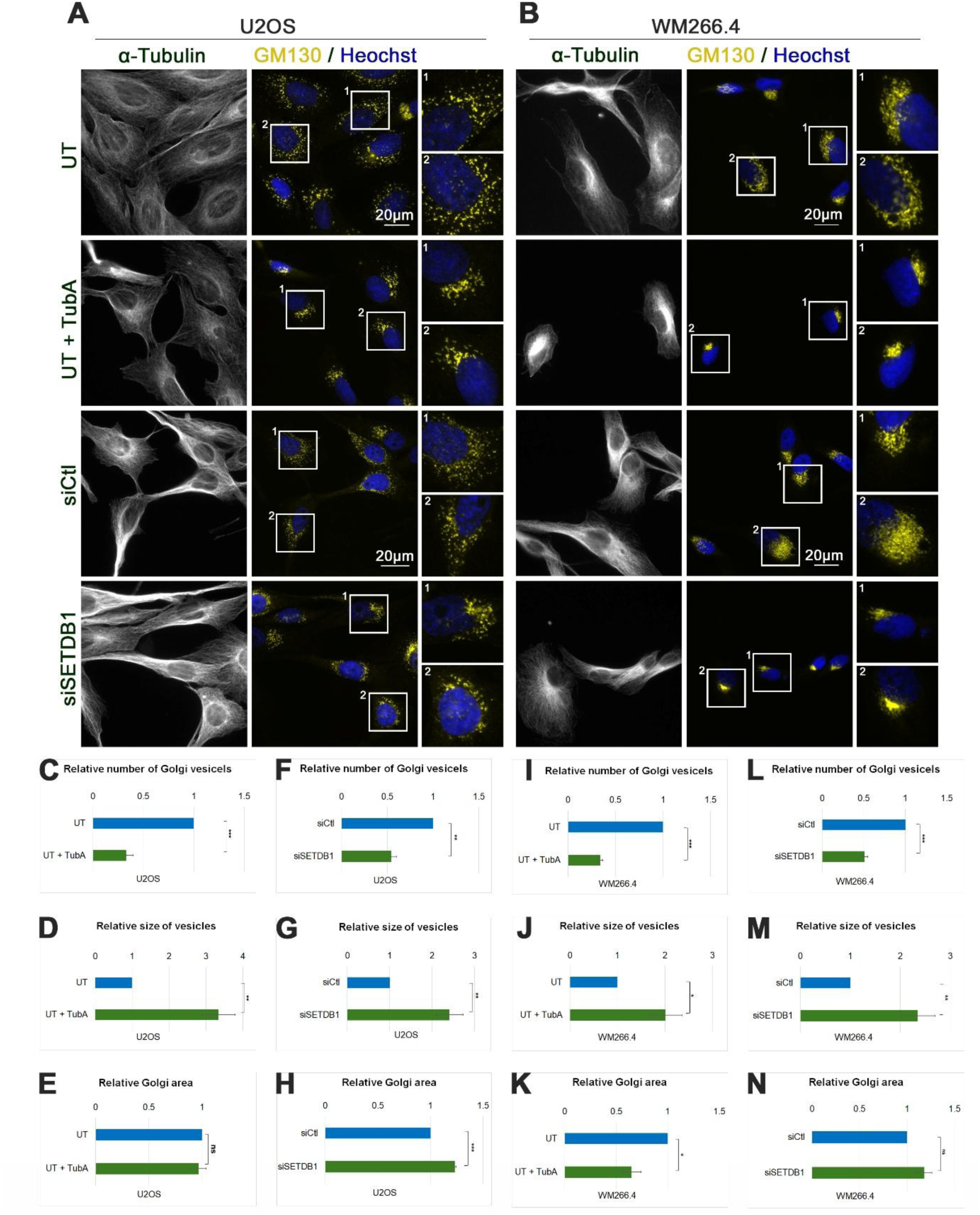
Knockdown of SETDB1 reduces Golgi fragmentation. **A-B.** U2OS cells (A) or WM266.4 cells (B) non-treated or treated for 1 h with 1.5 µM of HDAC6 inhibitor Tubastatin A (TubA), transfected with Ctl siRNA or SETDB1 siRNA immunostained for α-tubulin, the Golgi marker GM130. DNA was stained with Hoechst 33342. The Golgi outlined by white rectangles are magnified on the side of the micrographs. Scale bar: 20 μm. **C-N.** Quantification of the relative Golgi dispersal in U2OS cells (C-H) and WM266.4 cells (I-N). For each experiment, 21 cells per condition were measured by ImageJ using analyze particle tool for the number of Golgi vesicles per cell (C,F,I,L), average size of Golgi vesicle per cell (D,G,J,M) and total Golgi area (E,H,K,N). In non-transfected cells, in each experiment, the averages in TubA-treated cells were normalized to TubA-free cells. In siRNA-transfected cells, in each experiment, the averages in SETDB1 siRNA-transfected cells were normalized to Ctl siRNA-transfected cells. Average mean intensities from three independent experiments ± SE are presented. Statistical significance was calculated with Student’s t-test, \**p* < 0.05, \*\**p* < 0.01, \*\*\**p* < 0.001

## Discussion

HDAC6 is a key factor in regulation of MT dynamics and functions, mainly through its α-tubulin deacetylation activity. This deacetylation affects MT stability [11,12], intracellular transport, cell division and motility [59,61,62]. Thus, HDAC6 is connected to various pathologies including cancer and neurodegenerative diseases [21,22,63]. Here we identified a regulation mechanism of HDAC6 activity by SETDB1. SETDB1 co-immunoprecipitated with HDAC6 and promoted tubulin deacetylation both *in vitro* and *in vivo*. The ability of SETDB1 to attenuate the increase in tubulin acetylation upon paclitaxel treatment suggests that it may promote cancer cell resistance to paclitaxel treatment. A resistance that is a major challenge in cancer treatment with tubulin drugs [64]. Thus, targeting the SETDB1-HDAC6 axis could sensitize cancer cells to tubulin-targeting drugs.

The increase in SETDB1-dependent tubulin deacetylation by HDAC6 seems to occur on tubulin filaments rather than dimers: SETDB1 promoted the association of HDAC6 with MTs but not with tubulin dimers. This observation raises a mechanistic challenge; Lys40 of α-tubulin faces the MT lumen [51] while HDAC6 is not thought to be able to penetrate by itself into the MT lumen, unlike the tubulin acetyltransferase ATAT1 [10,16,18,19,52–54]. Still, HDAC6 could enter the MT lumen if lattice defects/vacancies are induced along the MT shaft. Damage along the shaft is repaired by incorporation of new GTP-bound tubulin dimers [45]. Notably, overexpression of SETDB1 led to increased GTP-αTubulin levels in MTs. Suggesting that SETDB1 could promote MT shaft damage to form penetration sites into the MT lumen for HDAC6 in a similar manner to the motor protein kinesin-1 [55].

To investigate the cellular consequences of the SETDB1-promoted HDAC6 activity we evaluated the morphology of the Golgi apparatus. Golgi dispersal is a very common phenomenon in cancer cells that is thought to support tumorigenesis by accelerating protein secretion and by altering the glycosylation pattern of membranal and secreted proteins [58,65,66]. Notably, knockdown of SETDB1 led to reduced dispersal of the Golgi in a couple of cancer cell types; U2OS and WM266.4. Interestingly, treatment of these cells with the HDAC6 inhibitor, TubA, led to a similar result. To verify this function of SETDB1 and the dependency of SETDB1 on HDAC6, we analyzed HeLa cells as well. In HeLa cells, in which the Golgi is organized in compact stacks, SETDB1 overexpression led to dispersal of the Golgi in an HDAC6-dependent manner. Both WT and CD forms of SETDB1 affected the Golgi organization. Moreover, overexpressed GFP-fused SETDB1 is mainly localized to the cytoplasm rather than the nucleus [41,67]. Thus, the ability of SETDB1 to affect the Golgi seems to rely on the cytoplasmic pool of SETDB1 that works in concert with HDAC6 in an independent manner of its methyltransferase activity. The difference in Golgi morphology is thought to be accompanied by alterations in the secretory pathway; different rates of protein processing in the secretory pathway as well as differences in the glycosylation pattern of the processed proteins. This can affect the interactions of tumor cells with both the extracellular matrix (ECM) and neighboring cells in the tumor microenvironment to promote tumor cells invasion and immune evasion [58,65,66]. Thus, as an oncogene and a MT-associated protein, SETDB1 could promote tumorigenesis by affecting both intracellular processes such as mitotic fidelity [41] and extracellular processes that are dependent on cancer cell interactions with the tumor microenvironment.

SETDB1 appears as part of a growing group of histone methylation factors with dual functions in regulation of chromatin organization and MT dynamics [23–28,31,33]. Currently, SETDB1 is unique in both its function and the mechanism it uses. While other methylation factors were found to stabilize MTs, SETDB1 promotes MT destabilization. Mechanistically, SETDB1 does so by promoting HDAC6 to deacetylate tubulin. By this mechanism SETDB1 may affect a broad range of cellular processes that are dependent on MTs and HDAC6 including organelle positioning, protein secretion, glycosylation, intercellular interactions and more.

## Supplementary Material

### Supplementary Figures

**Sup. Figure 1.**
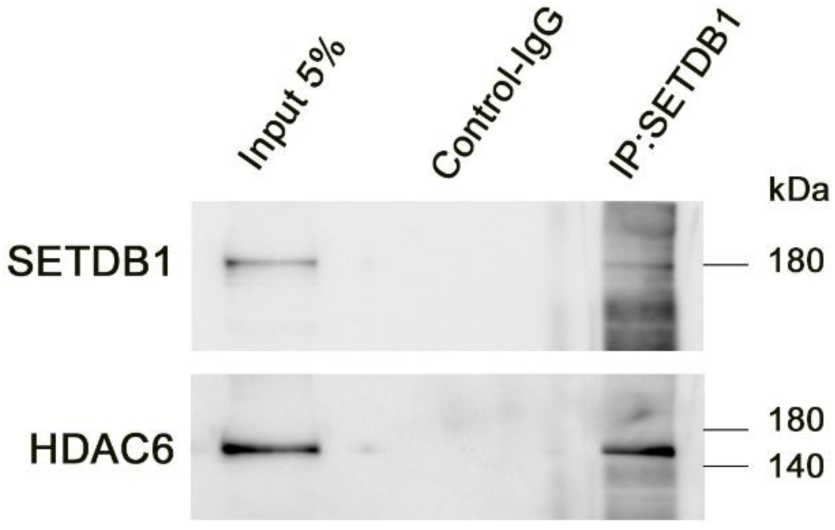
Co-immunoprecipitation of endogenous SETDB1 and HDAC6 in HEK293 cells. HEK293 cell lysates were immunoprecipitated with control rabbit IgG or an anti-SETDB1 antibody (SC-66884) and analyzed by Western blot for the indicated proteins.

**Sup. Figure 2.**
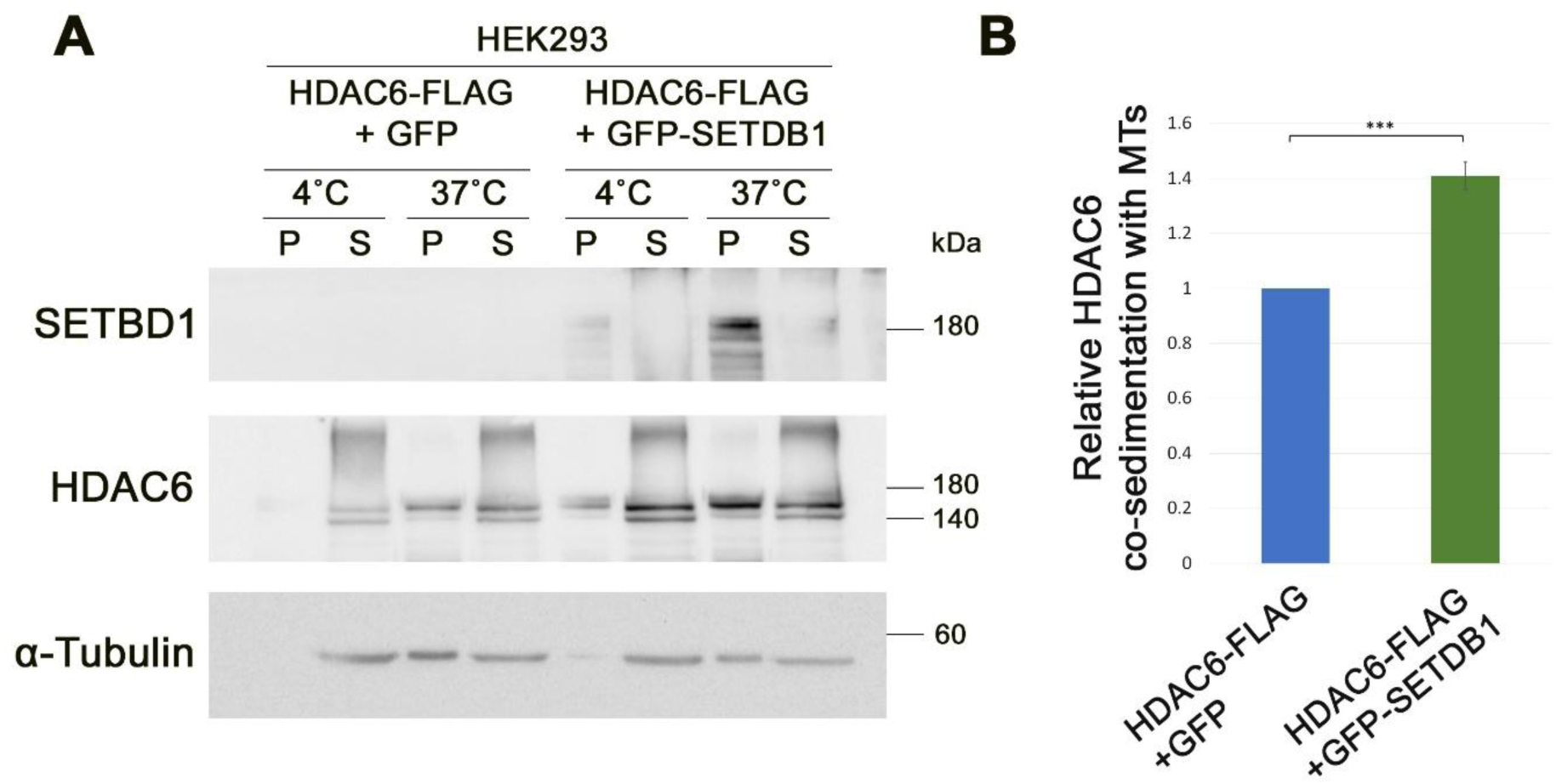
SETDB1 promotes HDAC6 co-sedimentation with MTs. **A.** MT co-sedimentation assay of HDAC6 in HEK293 cells. HEK293 cells overexpressing HDAC6-FLAG with GFP or GFP-SETDB1 used for MT co-sedimentation assay. The pellet (P) and supernatant (S) obtained after incubation at 37°C indicate MT-bound and unbound fractions, respectively. Incubation at 4°C serves as the MT-free control. **B.** The bar graphs show the relative levels of HDAC6 associated with MTs ± SE of four repetitions. In each repetition HDAC6 levels in the pellet were normalized to its levels at the supernatant at 37°C, and the ration in GFP expressing cells was set to 1. Statistical significance was calculated with Student’s *t*-test, \**P* < 0.05.

**Sup. Figure 3.**
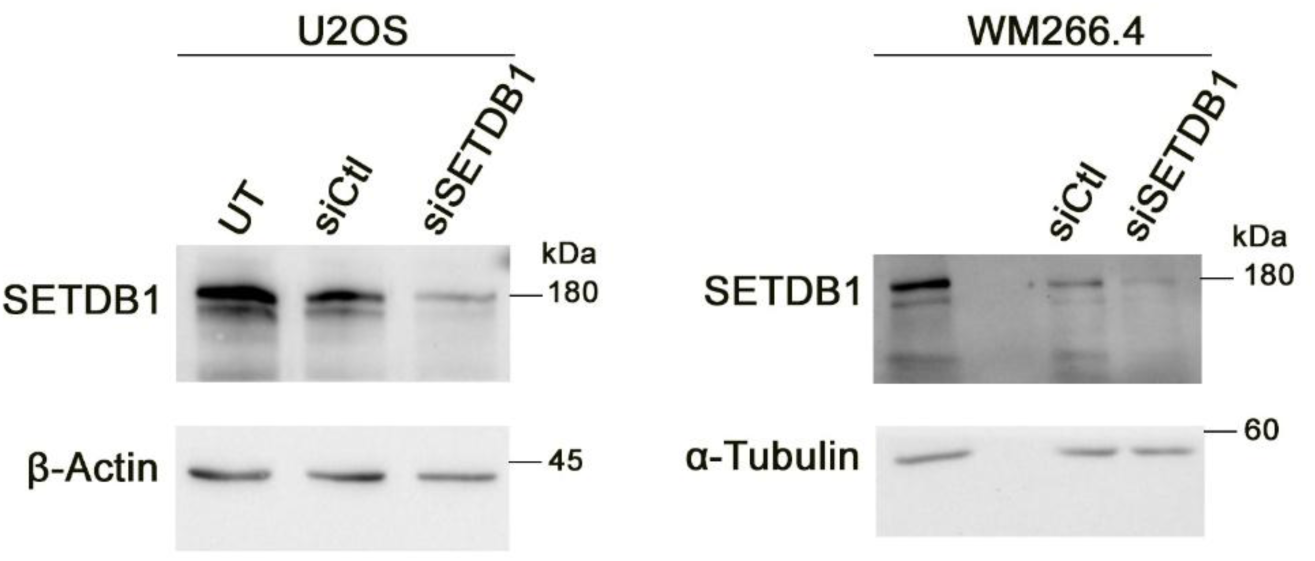
Knockdown of SETDB1. Western blot analysis of SETDB1 in untreated cells (UT), and cells transfected with Ctl siRNA or SETDB1 siRNA. B-Actin or a-Tubulin were used as a loading control.

