## Supplemental file for "SETDB1 promotes tubulin deacetylation and Golgi fragmentation by HDAC6"

Gowthaman Gunasekaran, Galia Gelman, Ohad Manshirov, Tamar Listovsky, and Gabi Gerlitz*
School of Biomedical Sciences, Faculty of Medicine, Ariel University, Ariel 40700, Israel.

### *Supplementary Material*

#### Supplementary Figures

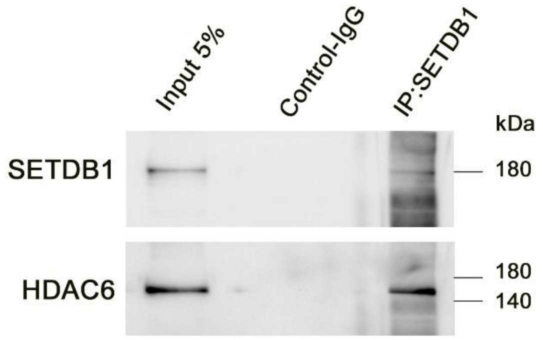


**Sup. Figure 1** Co-immunoprecipitation of endogenous SETDB1 and HDAC6 in HEK293 cells. HEK293 cell lysates were immunoprecipitated with control rabbit IgG or an anti-SETDB1 antibody (SC-66884) and analyzed by Western blot for the indicated proteins.

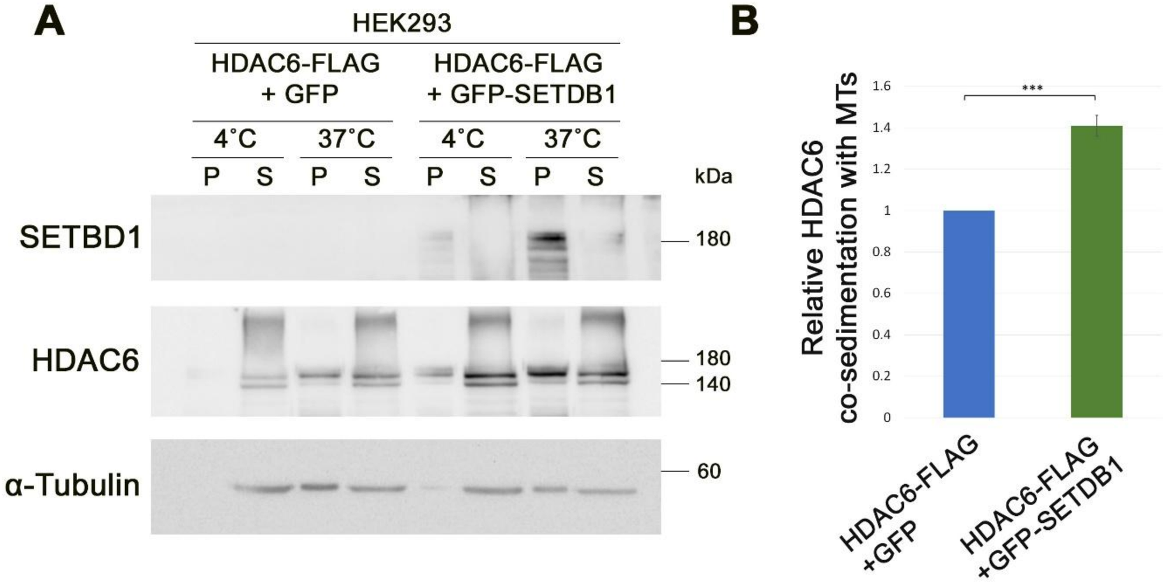


**Sup. Figure 2** SETDB1 promotes HDAC6 co-sedimentation with MTs. **A.** MT co-sedimentation assay of HDAC6 in HEK293 cells. HEK293 cells overexpressing HDAC6-FLAG with GFP or GFP-SETDB1 used for MT co-sedimentation assay. The pellet (P) and supernatant (S) obtained after incubation at 37ºC indicate MT-bound and unbound fractions, respectively. Incubation at 4ºC serves as the MT-free control. **B.** The bar graphs show the relative levels of HDAC6 associated with MTs ± SE of four repetitions. In each repetition HDAC6 levels in the pellet were normalized to its levels at the supernatant at 37ºC, and the ration in GFP expressing cells was set to 1. Statistical significance was calculated with Student's *t*-test, *$P < 0.05$.

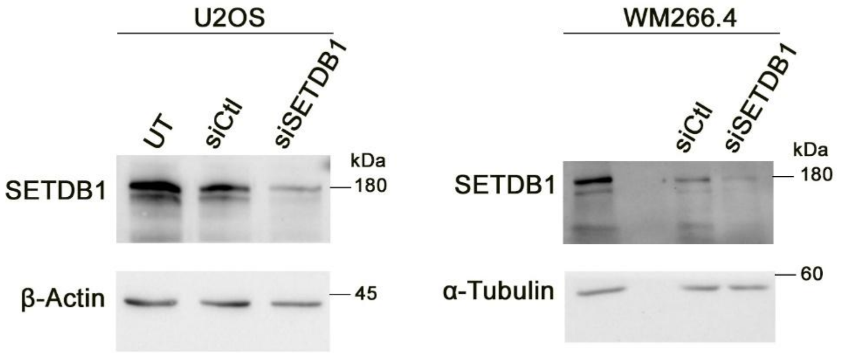


**Sup. Figure 3 Knockdown of SETDB1.** Western blot analysis of SETDB1 in untreated cells (UT), and cells transfected with Ctl siRNA or SETDB1 siRNA. B-Actin or a-Tubulin were used as a loading control.
